# White-nose syndrome restructures bat skin microbiomes

**DOI:** 10.1101/614842

**Authors:** Meghan Ange-Stark, Tina L. Cheng, Joseph R. Hoyt, Kate E. Langwig, Katy L. Parise, Winifred F. Frick, A. Marm Kilpatrick, Matthew D. MacManes, Jeffrey T. Foster

**Affiliations:** Department of Molecular, Cellular and Biomedical Sciences, University of New Hampshire, Durham, New Hampshire, 03824, USA; Department of Ecology and Evolutionary Biology, University of California, Santa Cruz, California 95064, USA; Bat Conservation International, Austin, Texas 78746, USA; Department of Biological Sciences, Virginia Tech, Blacksburg, Virginia 24061, USA; Pathogen and Microbiome Institute, Northern Arizona University, Flagstaff, Arizona 86011, USA

**Keywords:** bat populations, disease ecology, microbiome, *Myotis lucifugus*, *Myotis septentrionalis*, *Perimyotis subflavus*, *Pseudogymnoascus destructans*, white-nose syndrome

## Abstract

The skin microbiome is an essential line of host defense against pathogens, yet our understanding of microbial communities and how they change when hosts become infected is limited. We investigated skin microbial composition in three North American bat species (*Myotis lucifugus*, *Eptesicus fuscus*, and *Perimyotis subflavus*) that have been impacted by the infectious disease, white-nose syndrome, caused by an invasive fungal pathogen, *Pseudogymnoascus destructans*. We compared bacterial and fungal composition from 154 skin swab samples and 70 environmental samples using a targeted 16S rRNA and ITS amplicon approach. We found that for *M. lucifugus*, a species that experiences high mortality from white-nose syndrome, bacterial microbiome diversity was dramatically lower when *P. destructans* is present. Key bacterial families—including those potentially involved in pathogen defense—significantly differed in abundance in bats infected with *P. destructans* compared to uninfected bats. However, skin bacterial diversity was not lower in *E. fuscus* or *P. subflavus* when *P. destructans* was present, despite populations of the latter species declining sharply from white-nose syndrome. The fungal species present on bats substantially overlapped with the fungal taxa present in the environment at the site where the bat was sampled, but fungal community composition was unaffected by the presence of *P. destructans* for any of the three bat species. This species-specific alteration in bat skin bacterial microbiomes after pathogen invasion may suggest a mechanism for the severity of WNS in *M. lucifugus*, but not for other bat species impacted by white-nose syndrome.

## Introduction

The microbiome is defined as the collection of microbes (composed of bacteria, bacteriophage, fungi, protozoa, and viruses) that live in and on an organism (Turnbaugh et al., 2006). Microbiomes are increasingly being recognized as critical components of host health, directly influencing a range of biochemical and physiological processes (Cho & Blaser, 2012). For mammalian skin microbiomes in particular, specific taxa are recognized as common inhabitants although these communities are just beginning to be characterized (Ross, Müller, Weese, & Neufeld, 2018). Researchers have identified several bacterial species that are associated with skin disease in humans (Findley & Grice, 2014; Kong et al., 2012; Zeeuwen, Kleerebezem, Timmerman, & Schalkwijk, 2013), including *Staphylococcus aureus*, which is linked to atopic dermatitis in children (Kong, 2011), *Corynebacterium minutissimum*, the agent of erythrasma, a chronic, superficial infection that causes lesions, and *Streptococcus pyogenes*, the most common agent of cellulitis, a diffuse inflammation of loose connective tissue (Aly, 1996). In wildlife, a pathogenic microbe, *Batrachochytrium dendrobatidis*, the fungal causative agent of chytridiomycosis in the skin of amphibians, has devastated amphibian populations worldwide (Berger et al., 1998; Longcore, Pessier, & Nichols, 1999; Piotrowski, Annis, & Longcore, 2004). However, many species of bacteria exist as commensals and confer benefits to their hosts including defense against pathogens, metabolism, and reproduction (Cho & Blaser, 2012).Examples include various staphylococcal species that inhibit skin inflammation after injury (Lai et al., 2009), as well as *Staphylococcus epidermidis*, which has been known to protect humans from an array of pathogenic bacteria such as *S. aureus*. Likewise, cutaneous microbes on amphibians might even play a protective role, allowing for resistance to pathogenic fungi (Bataille, Lee-Cruz, Tripathi, Kim, & Waldman, 2016; Belden & Harris, 2007; Harris, James, Lauer, Simon, & Patel, 2006; Woodhams et al., 2007).

While numerous microbiome studies have been conducted for wildlife diseases and correlations between pathogen colonization and microbial diversity have been observed, determining causality remains challenging. Previous studies have shown that pathogens can alter the microbiome. For example, *B. dendrobatidis* (Bd) disturbs the frog skin microbiome during both natural and experimental infection; bacterial richness was significantly lower on Bd-infected frogs compared with uninfected frogs (Jani & Briggs, 2014). Similarly, snake fungal disease was correlated with a reduction in bacterial and fungal diversity in an endangered rattlesnake (Allender, Baker, Britton, & Kent, 2018). In Sea Star Wasting Disease, changes in microbial community composition occur during disease progression, with decreasing species richness in the late stages of the disease (Lloyd & Pespeni, 2018). With such perturbations to microbiomes following the introduction of various pathogens, continued investigation of skin microbial inhabitants—both pathogens and commensals—is crucial to understanding microbial pathogenesis and the role of the microbiome in animal health.

White-nose syndrome (WNS) in bats, caused by the fungal pathogen *Pseudogymnoascus destructans* (Pd), has become another prominent example of a lethal skin infection in wildlife (Blehert et al., 2009; Lorch et al., 2011; Warnecke et al., 2012). WNS is characterized by cutaneous infection during hibernation (Meteyer et al., 2009). The onset and growth of Pd on bat skin causes dehydration, fat loss, and electrolyte imbalance that disrupts bats’ natural torpor cycle resulting in the depletion of fat reserves and mortality (Verant et al., 2014; Warnecke et al., 2013). WNS emerged in North American bats in winter 2005/2006 (Blehert et al., 2009; Lorch et al., 2011; Warnecke et al., 2012) and has caused massive declines in bat populations throughout its spread across the Northeast and Midwestern US and Eastern Canada (Frick et al., 2010, 2015; Langwig et al., 2012, 2017; Langwig, Hoyt, et al., 2015). Population declines at hibernating sites have been severe, leading to local extirpation of some species (Frick et al., 2015; Langwig et al., 2012), and much smaller persisting populations of other species (Frick et al., 2017; Langwig et al., 2017; Reichard et al., 2014). Once surviving bats leave hibernacula, increases in body temperature and restored immune function enable bats to clear infection (Langwig, Frick, et al., 2015; Meteyer et al., 2011). However, Pd conidia remaining in caves can persist in the absence of bats, resulting in reinfection the following winter (Hoyt et al., 2015; Langwig, Frick, et al., 2015; Lorch et al., 2013).

Mortality rates from WNS vary among bat species (Langwig et al., 2012, 2016), and environmental conditions of hibernacula may be strong predictors of species impacts (Langwig et al., 2016). However, additional variation that could not be explained by environmental conditions alone suggests that interactions with other processes (e.g. behavioral, immune response, and microbiomes), may play a role in WNS susceptibility (Langwig et al., 2016). While investigations into the microbiomes of North American bats have been conducted to help characterize their skin microbiota (Avena et al., 2016; Lemieux-Labonté, Simard, Willis, & Lapointe, 2017; Winter et al., 2017), host-microbial interactions and their impact on bat health remains an important knowledge gap (Avena et al., 2016). Comparing the bacterial and fungal microbial composition of Pd-positive and Pd-negative bats across the range of Pd spread in North America provides an opportunity to investigate the changes a pathogenic fungus has on the host skin microbial community across several bat species. In addition, distinguishing resident bat skin microbiota from microbiota found in the surrounding environment can further contribute to our understanding of how host skin microbiomes are shaped and interact with local microbiota (Cogen, Nizet, & Gallo, 2008).

We examined the epidermal microbiomes of three North American bat species: *Myotis lucifugus*, *Perimyotis subflavus*, and *Eptesicus fuscus*. These species were selected because they differ in both sociality and susceptibility to WNS and have been well sampled across a broad range. Both *M. lucifugus* and *P. subflavus* are heavily impacted by WNS (Frick et al., 2017, 2015; Langwig et al., 2012), while *E. fuscus* is less affected (Langwig et al., 2012, 2016). We used a targeted 16S rRNA and ITS amplicon approach comparing Pd-positive and Pd-negative bat skin swabs to examine interactions between bat epidermal bacterial and fungal microbiomes. We also classified the resident and transient microbes by comparing bat skin swabs to environmental (substrate) swabs to determine which members of the microbiome potentially serve as commensals and which may be transients from the environment. We hypothesized that bats infected with Pd will have lower skin microbial diversity but only for species that are heavily impacted by WNS (*M. lucifugus* and *P. subflavus*; not *E. fuscus*), and bacterial species that occur in higher abundance on Pd-positive bats will enhance the microbial communities’ anti-fungal properties. Additionally, we predicted that we would find a significant difference in the microbiome composition between bat skin swabs and environmental swabs because only a fraction of bacteria and fungi in the environment will be able to colonize bats’ skin.

## Methods

### Data collection

Three species of hibernating bats (*E. fuscus*, big brown bat; *P. subflavus*, tri-colored bat; and *M. lucifugus*, little brown bat) were sampled at 34 hibernacula in 22 states throughout the eastern U.S. (Fig. 1). Samples were collected using epidermal swabbing of bats during winter hibernacula surveys conducted November–March of six consecutive winters from 2009/2010 to 2015/2016. Participating biologists were provided a detailed sampling protocol and video instructions to standardize sampling across sites and years (Frick et al., 2017). Epidermal swab samples were collected by dipping a sterile polyester swab in sterile water and rubbing the swab five times over the bat’s forearm and muzzle (Langwig, Frick, et al., 2015). Substrate swab samples were simultaneously collected from the ceiling or walls of each hibernaculum at least 10 cm from a roosting bat. After collection, swabs were placed in vials containing RNAlater (Thermo Fisher Scientific, Waltham, MA) and were subsequently stored at −20 °C, until DNA extraction.

**Figure 1.**
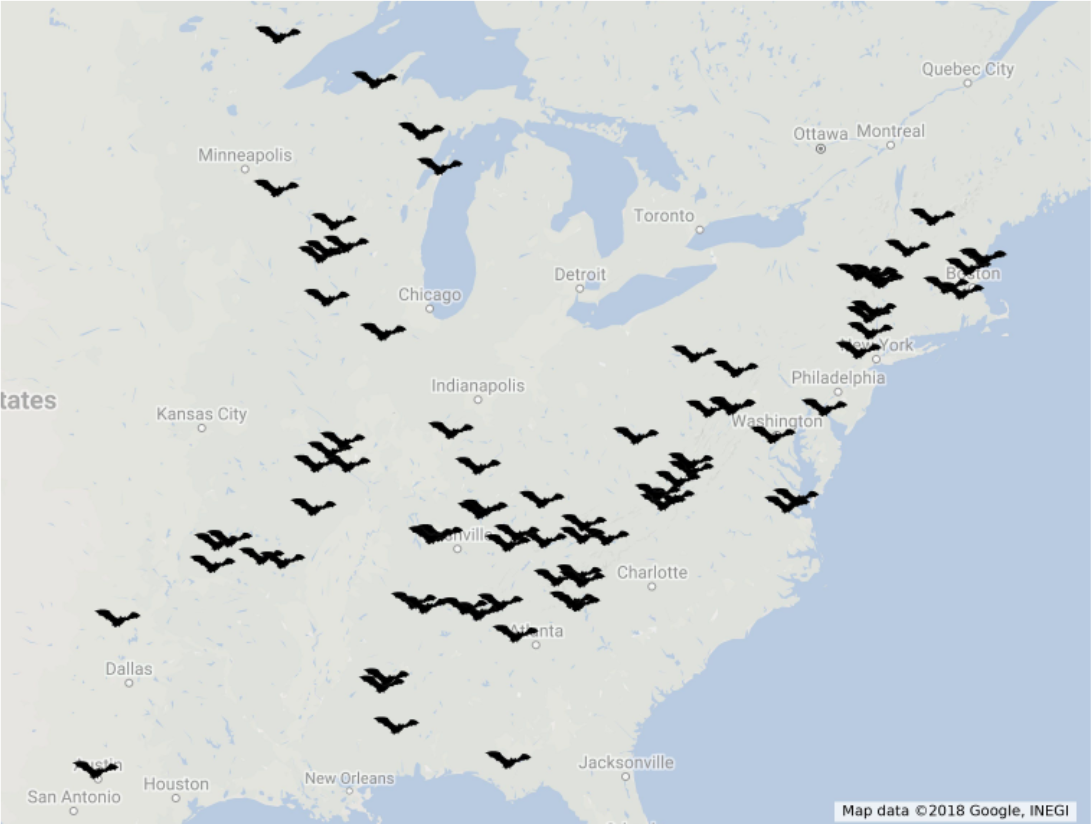
Sample collection distribution map. Bat and substrate samples were collected from 34 hibernacula in 22 of the 33 United States confirmed with presence of *Pseudogymnoascus destructans* from winters 2009/2010 to 2015/2016.

### DNA extraction and testing

Bat and substrate swabs were extracted using DNeasy Blood and Tissue extraction kits (Qiagen, Valencia, CA). The protocol was modified for fungal extractions to include lyticase during the lysis step in addition to the proteinase K and buffer ATL (Shuey, Drees, Lindner, Keim, & Foster, 2014). Pd DNA quantity for each sample was tested using a quantitative PCR assay (Muller et al., 2013). Samples were considered WNS positive if at least one of two qPCR cycle thresholds (Ct) was below 40.

### Amplification and sequencing

The DNA extracted from samples was prepped for sequencing following the Earth Microbiome Project 16S Illumina Amplicon Protocol and ITS Illumina Amplicon Protocol (earthmicrobiome.org; (Caporaso et al., 2012)) to test for bacterial and fungal taxa, respectively. Completed 16S rRNA and ITS libraries were submitted for 2 × 250 bp paired-end sequencing on a HiSeq 2500 at the University of New Hampshire Hubbard Center for Genome Studies.

16S libraries were prepared by first amplifying the hypervariable region V4 of the 16S small subunit ribosomal gene with forward (barcoded) primer 515FB and reverse primer 806RB and an annealing temperature of 50 °C for 60 s. Negative controls were included during amplification to account for possible contamination. Amplicons from each sample were run on an agarose gel to verify presence of PCR product, with an expected band size for 515f-806r of ∼300–350 bp. Amplicons were then quantified via a Qubit fluorometer (Thermo Fisher Scientific). An equal amount of amplicon from each sample (240 ng) was combined into a single, sterile tube. Amplicon pools were then cleaned using a QIAquick PCR Purification Kit (Qiagen), following the manufacturer’s instructions. An aliquot of the final pool was submitted for sequencing with the 16S forward and reverse sequencing primers as well as the index sequencing primer.

ITS libraries were prepared by first amplifying the nuclear ribosomal internal transcribed spacer (ITS) region with the forward primer ITS1f and reverse (barcoded) primer ITS2 using an annealing temperature of 52 °C for 30 s. Negative controls were included during amplification to account for possible contamination. Amplicons from each sample were run on an agarose gel to verify presence of PCR product, with an expected band size for ITS1f-ITS2 of ∼230 bp. Amplicon cleanup and pooling followed the procedures detailed above for 16S rRNA libraries.

### Data processing

Adapter sequences were removed using cutadapt (Martin, 2011). Sequence quality control, feature table construction, and raw sequence merging were performed using DADA2 version 1.8 in QIIME 2 (https://qiime2.org/). Unpaired reads were removed from the dataset. Feature tables for each sequence type contained counts (frequencies) of each unique sequence in each sample within the dataset. Additionally, this step detected and corrected Illumina amplicon sequence data, while also filtering any phiX reads as well as chimeric sequences. Best practices for microbial community analyses require that contamination caused by reagent and laboratory steps be removed as their presence can critically impact sequence-based microbial analyses (Knight et al., 2018; Salter et al., 2014). In our specific work, 16S sequence contamination present as a by-product of lyticase use during the fungal DNA extraction step was filtered out by removing reads corresponding to *Arthrobacter luteus* and related species (i.e.. *Cellulosimicrobium* spp.). In order to conduct diversity analyses, a multiple sequence alignment was constructed to create an aligned sequences feature table using mafft version 7.407 (https://mafft.cbrc.jp/alignment/software/; (Katoh & Standley, 2013)). Highly variable positions were removed to reduce noise in the phylogenetic tree. FastTree version 2.1 (http://www.microbesonline.org/fasttree/; ((Price, Dehal, & Arkin, 2010)) was applied to the filtered alignment, creating an unrooted phylogenetic tree where midpoint rooting was done at the midpoint of the longest tip-to-tip distance to create a rooted tree.

#### Microbial diversity analyses

QIIME2 version 2018.2 was used to conduct diversity analyses with accompanying statistical tests. The core metrics phylogenetic method, which rarefied the feature table to a user-specified depth, computed and provided interactive visualizations for alpha and beta diversity metrics, while also generating principal coordinate analysis plots (PCoA) plots using Emperor (https://biocore.github.io/emperor/) for beta diversity analyses.

Associations between categorical metadata (Pd infection status, sample type, phylogeny, etc.) and alpha diversity data were conducted to determine any significant differences between metadata groups in QIIME2. Likewise, sample composition in the context of categorial metadata using beta diversity metrics was analyzed using PERMANOVA (Anderson & Walsh, 2013). These tests determined which specific pairs of metadata groups differed from one another.

#### Linear mixed-effects modeling analyses

To better understand the variables potentially driving skin bacterial diversity differences between Pd-positive and Pd-negative bats, we fit a linear mixed effects model that included Pd presence (coded as 0/1), date, site latitude, and monthly average outside temperature of sample collection site as fixed effects and sample collection site as a random effect to account for multiple bats sampled at each site. Pd presence was determined using quantitative PCR (Muller et al., 2013) as described above. Monthly average temperatures of the sample collection date were collected from the National Oceanic and Atmospheric Association Climate Database (https://www.ncdc.noaa.gov/). Latitude at the sample collection site was included as a covariate to account for potential latitudinal diversity gradients in terrestrial bacteria (Andam et al., 2016), while winter sample selection was done to account for seasonal effects on microbial composition. Models were fit using the lmer function in the R package lme4. We assumed a Gaussian distribution and checked model residuals to confirm normality for each response variable (richness, evenness, and Shannon index) for each species.

#### Taxonomic identification and differential abundance analyses

Identifying the taxonomic composition of 16S rRNA sequences required the use of a pre-trained Naive Bayes classifier as well as the QIIME2 feature classifier plugin, which was trained on the Greengenes 13_8 99% OTUs where the sequences have been trimmed to only include 250 bases from the region of the 16S rRNA that was sequenced in the analysis (the V4 region, bound by the 515F/806R pair). ITS taxonomic identification leveraged the UNITE database version 7.2 (https://unite.ut.ee/). The linear discriminant analysis (LDA) effect size (LEfSe) was used to conduct microbial differential abundance analyses (http://huttenhower.sph.harvard.edu/lefse/; (Segata et al., 2011)).

## Results

For bacteria, 3,663,044 16S rRNA sequences amplified in 224 samples. Of those 224 samples, 154 were bat skin swabs and the remaining 70 samples were substrate swabs. A mean frequency of 12,162 16S rRNA sequences were obtained per bat skin sample, ranging from 1,801 to 381,143 reads. A mean frequency of 25,572 16S sequences were obtained per substrate sample, ranging from 1,003 to 391,366 reads. For fungi, 17,257,384 ITS sequences amplified from 498 samples. Of those 498 samples, 375 samples were bat skin swabs and the remaining 123 were substrate swabs. A mean of 37,092 ITS sequences were obtained per bat skin sample, ranging from 1,013 to 480,261 reads. A mean of 27,217 ITS sequences were obtained per substrate sample, ranging from 1,019 to 273,092 reads. Raw sequencing reads are available at the NCBI Short Read Archive (accession number: PRJNA533244).

### Bacterial and fungal skin microbiome dissimilarities between Pd-negative E. fuscus, M. lucifugus, and P. subflavus

Jaccard distance matrices that quantitatively measured bacterial community dissimilarity showed that—when measuring bacterial species presence and absence among bat species—*E. fuscus* and *P. subflavus* were the most similar in bacterial community composition (Fig. S3; *p* = 0.19), while *E. fuscus* and *M. lucifugus* were the most dissimilar in bacterial community composition (Fig. S3, *p* = 0.01). Jaccard distance matrices for fungal community dissimilarity showed high levels of fungal community dissimilarity between all species pairs (Fig. S4; *E. fuscus* and *M. lucifugus*, *p* = 0.001; *E. fuscus* and *P. subflavus*, *p* = 0.001; *M. lucifugus* and *P. subflavus*, *p* = 0.001).

### Characterization of the Pd-negative bacterial and fungal microbiomes of E. fuscus, M. lucifugus, and P. subflavus

The bacterial (Fig. S1) and fungal (Fig. S2) skin microbiomes of Pd-negative *E. fuscus*, *M. lucifugus*, and *P. subflavus* were characterized to better understand the composition of skin microbiota before WNS invasion. These samples were all taken from bats from areas where WNS and Pd had not yet been detected (i.e., at least one year prior to Pd arrival). The three most common bacterial orders across all three bat species were *Pseudomonadales*, *Enterobacteriales*, and *Actinomycetales* but the relative abundance and dominance of each order differed among bat species (Fig. 2). Bacterial communities on *E. fuscus* (Fig. 2, A1) were dominated by a single order, whereas skin microbiomes were much more even on *M. lucifugus* (Fig. 2, B1) and *P. subflavus* (Fig. 2, C1). *Pseudomonadales* was the most common bacterial order for both *E. fuscus* (59.5% of OTUs), and *P. subflavus* (20.9%), and the third most common bacteria on *M. lucifugus* (12.0%). *Actinomycetales* was also a dominant order on all three species, making up 6.5%, 13.5%, and 11.0% of bacterial OTUs on *E. fuscus, M. lucifugus*, and *P. subflavus*, respectively.

**Figure 2.**
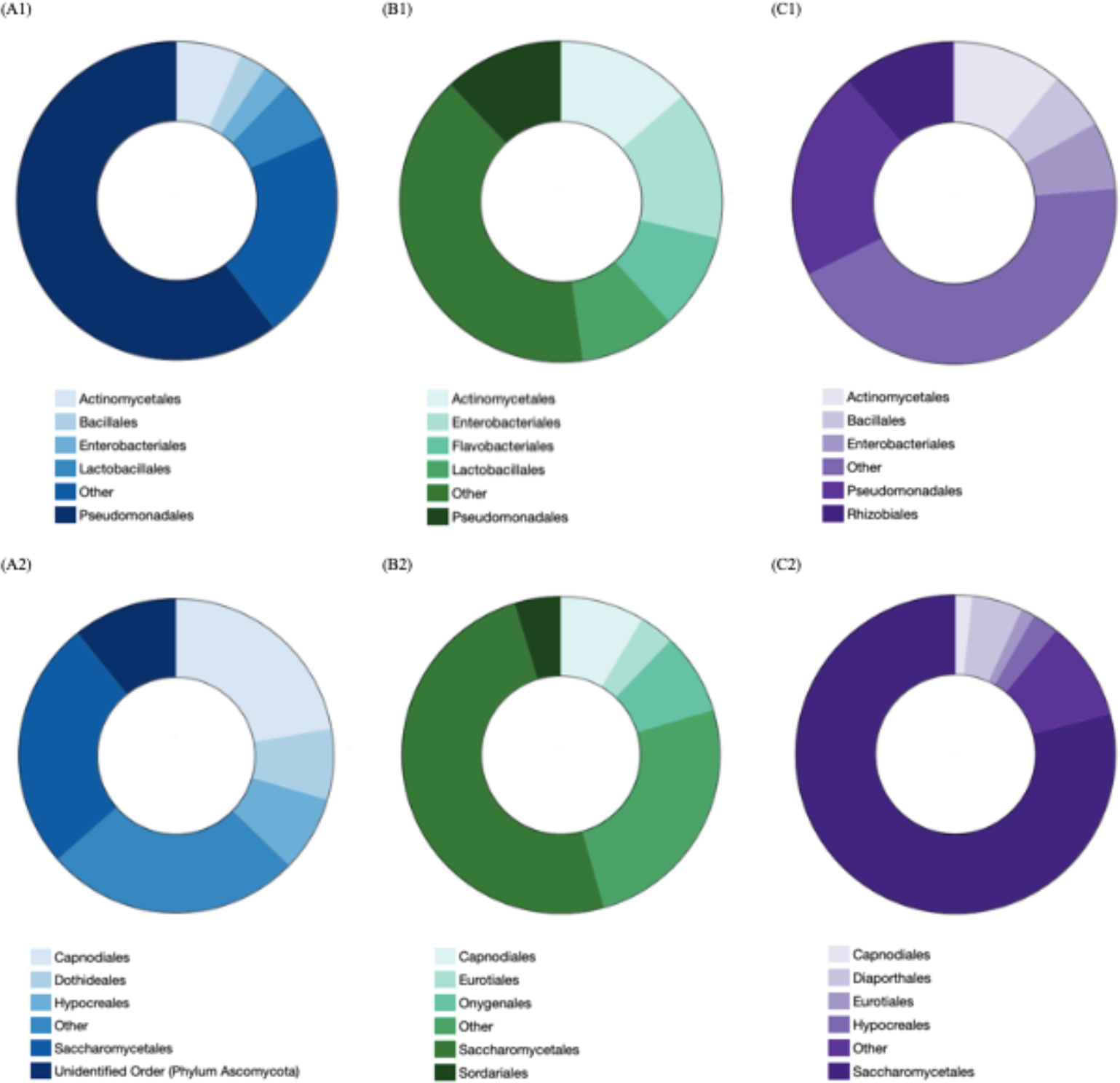
Bacterial and fungal microbiome taxonomy characterization. Bacterial characterization at the order level showed that for *Eptesicus fuscus* (A1) 59.52% of the bacterial OTUs were assigned to the order *Pseudomonadales*, for *Myotis lucifugus* (B1), *Enterobacteriales* was he most abundant bacterial order with 15.05% of the bacterial OTUs, and *Pseudomonadales* was the most abundant bacterial order for *Perimyotis subflavus* (C1) with 20.93% of the bacterial OTUs. Fungal microbiome characterization at the order level showed that for all three bat species, *Saccharomycetales* was the most abundant fungal order; however, it’s abundance varied significantly between each species (*E. fuscus*, 25.54%; *M. lucifugus*, 49.41%; *P. subflavus*, 78.91%).

*Saccharomycetales* was the most common fungal order across all three bat species; however, its relative abundance varied by bat species. *Eptesicus fuscus* (Fig. 2, A2) displayed higher levels of evenness with relatively similar abundances of *Saccharomycetales* and its second most abundant order, *Capnodiales*, while fungal communities on *M. lucifugus* (Fig. 2, B2) and *P. subflavus* (Fig. 2, C2) were both dominated by *Saccharomycetales*. *Capnodiales* and *Onygenales*—the second and third most abundant fungal orders for *M. lucifugus*, respectively—each made up less than 10% of its fungal microbiome. The same was true for *P. subflavus*, with *Diaporthales* and *Hypocreales* each making up less than 5% of the fungal microbiome as the second and third most abundant fungal orders.

### Microbial diversity analyses: microbiome differences between Pd-positive and Pd-negative bats

Species richness, evenness and Shannon diversity were similar in Pd-positive and Pd-negative *E. fuscus* bat samples for both bacterial (Fig. 3, A1; Evenness: *χ*^2^ = 0.69, *p* = 0.72, Richness: *χ*^2^ = 0.35, *p* = 0.37, and Shannon Diversity: *χ*^2^ = 0.60, *p* = 0.62), and fungal microbiomes (Fig. 3, B1; Evenness: *χ*^2^ = 0.33, *p* = 0.34, Richness: *χ*^2^ = 0.16, *p* = 0.17, and Shannon Diversity: *χ*^2^ = 0.20, *p* = 0.21). Fungal communities for *M. lucifugus* were also similar in species diversity (Fig. 3, B2; Richness: *χ*^2^ = 0.47, *p* = 0.47, Shannon Diversity: *χ*^2^ = 0.90, *p* = 0.91) and evenness (Fig. 3, B2; *χ*^2^ = 0.99, *p* = 1.00) when comparing Pd-negative and positive samples. In contrast, bacterial microbiomes in *M. lucifugus*, were less diverse (Fig. 3, A2; Richness: *χ*^2^ = 0.013, *p* = 0.013, Shannon Diversity: *χ*^2^ = 0.002, *p* = 0.002) and less even (Fig. 3, A2; *χ*^2^ = 0.013, *p* = 0.014) in Pd positive than negative samples. Bacterial microbiomes for *P. subflavus* were similar between Pd-positive and negative bats (Fig. 3, A3; Evenness: *χ*^2^ = 0.10, *p* = 0.10, Richness: *χ*^2^ = 0.19, *p* = 0.20, and Shannon Diversity: *χ*^2^ = 0.10, *p* = 0.10), whereas fungal microbiomes had higher evenness (Fig. 3, B3; *χ*^2^ = 0.001, *p* = 0.001) and diversity (Fig. 3, B3; *χ*^2^ = 0.002, *p* = 0.002) in the Pd-positive group than the Pd-negative group, but not in richness (Fig. 3, B3; *χ*^2^ = 0.26, *p =* 0.26). Thus, there appeared to be substantial variation in the effects of the presence of Pd on bat microbiomes and that variation appears to differ by bat host.

**Figure 3.**
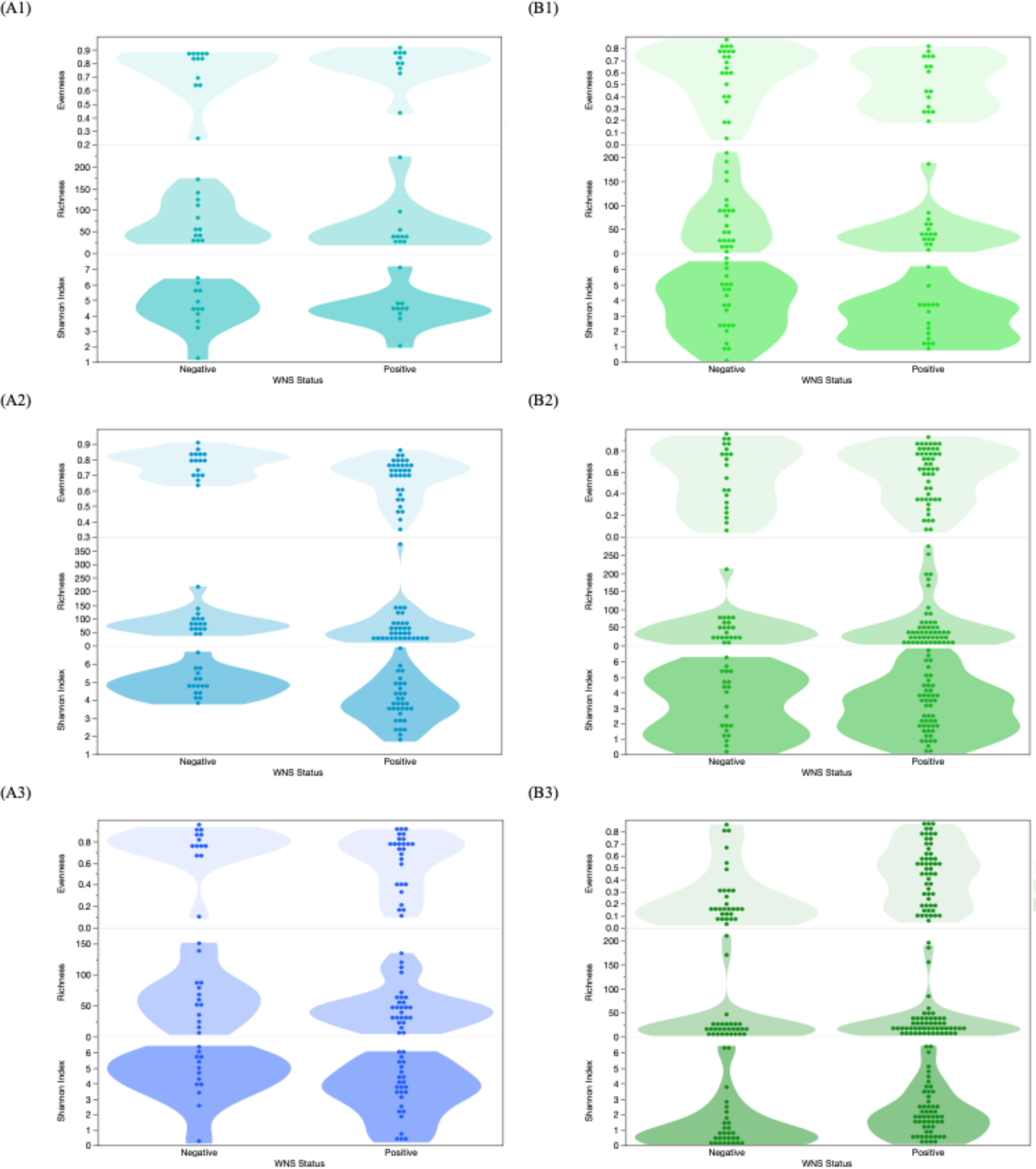
Measures of bacterial and fungal evenness, richness, and Shannon indices between Pd-positive and Pd-negative bats. There were no significant differences when comparing (A1) bacterial or (B1) fungal diversity between Pd-positive and Pd-negative *Eptesicus fuscus*. (A2) Bacterial diversity comparisons in *Myotis lucifugus* found the evenness, richness and Shannon diversity were all significantly higher in Pd-negative bats. (B2) Fungal diversity comparisons in *M. lucifugus* did not indicate any difference between Pd-positive and Pd-negative bats. (A3) There were no significant differences when comparing bacterial diversity between Pd-positive and Pd-negative *Perimyotis subflavus*; however, (B3) fungal diversity results from *P. subflavu*s indicate that Pd-positive bats have an higher fungal evenness and Shannon diversity.

### Does Pd influence M. lucifugus’ skin bacterial microbiome?

We found that fungal diversity for *M. lucifugus* was most influenced by both average temperature and latitude, as one or both of these covariates were significant for all three response variables (Richness: monthly average temperature, *p* = 0.0424, PE = 1.4, SE = 0.66; Evenness: monthly average temperature, *p* = 0.0241, PE = 0.008, SE = 0.003, and latitude, *p* = 0.0314, PE = 0.03, SE = 0.01; Shannon Diversity: monthly average temperature, *p* = 0.0046, PE = 0.07, SE = 0.02, and latitude, *p* = 0.0456, PE = 0.18, SE = 0.24). In contrast, results for *P. subflavus* indicated that sample date most significantly influenced fungal richness (*p* = 0.01, PE = 0.26, SE = 0.09) and Shannon diversity (*p* = 0.0421, PE = 0.008, SE = 0.004) when compared to the other covariates. We found no clear effect of Pd presence, temperature, latitude, and sampling date on bacterial evenness or diversity for *P. subflavus* or *E. fuscus*; however, Pd presence (0/1) on *M. lucifugus* significantly decreased bacterial diversity (Evenness: *p* = 0.012, PE = 0.05, SE = 0.02; Shannon Diversity: *p* = 0.0009, PE = 0.66, SE = 0.18). Increases in Pd load in a sample decreased Shannon diversity in *M. lucifugus*, although the result was not statistically significant (Fig. S5).

### Taxonomic differential abundance analyses in Pd-positive and Pd-negative M. lucifugus

Comparisons of of skin microbial diversity between Pd-positive and Pd-negative *M. lucifugus* uncovered substantial differences in bacterial taxa between the two groups, as both richness and evenness were significantly lower in the Pd-positive bats. A differential abundance analysis (Fig. 4) indicated that one family of bacteria, *Pseudonocardiaceae*, was overly abundant among Pd-positive *M. lucifugus*. Conversely, bacterial families *Cytophagaceae* and *Rhizobiaceae*, as well as bacterial order *Chromatiales*, were abundant in Pd-negative *M. lucifugus*.

**Figure 4.**
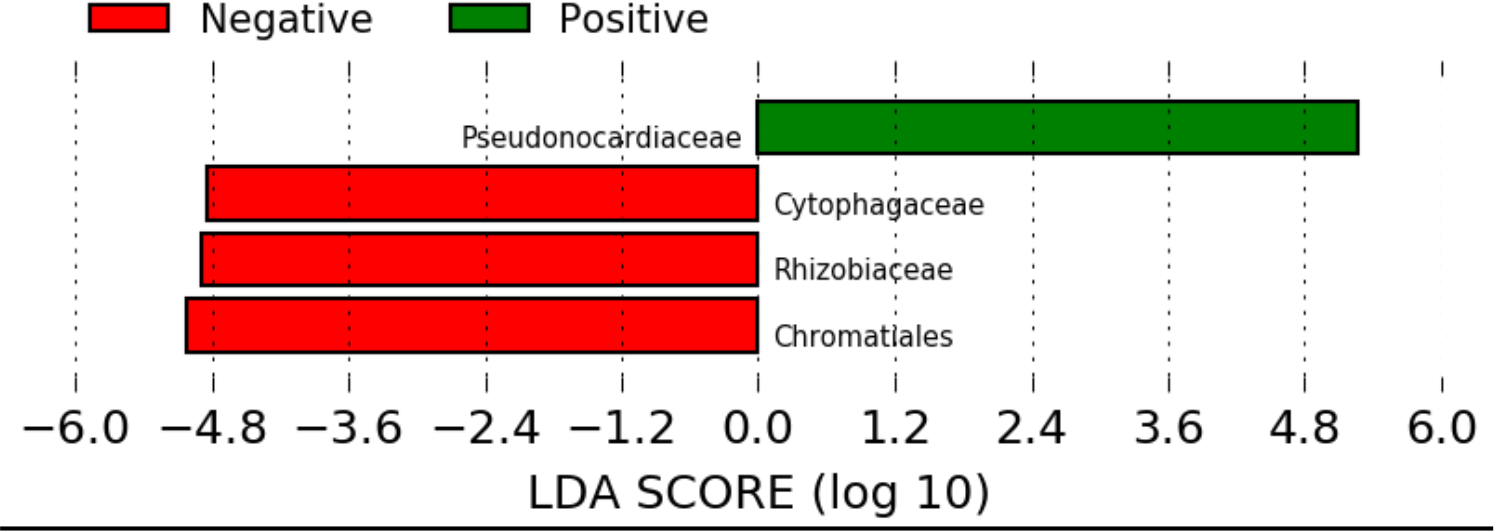
Relative abundances of bacterial epidermal communities of Pd-positive and Pd-negative *Myotis lucifugus*. Measures of relative abundance between the bacterial epidermal communities from swabs taken from Pd-positive and Pd-negative *M. lucifugus* revealed an overabundance of the bacterial family *Pseudonocardiaceae* in Pd-positive bats, while unaffected bats showed an overabundance of bacterial families *Cytophagaceae* and *Rhizobiaceae*, as well as the bacterial order *Chromatiales*.

### Comparison of skin microbiome between bats and substrates

Bacterial diversity was significantly higher on substrates than on bats (Fig. 5A, *χ*^2^ = 0.0001, p <0.0001) with 17 overly abundant bacterial families represented in the combined substrate samples and only four overly abundant bacterial families represented in combined bat samples (Fig. S6). Fungal communities between the two sample types, however, were similar in diversity (Fig. 5B, *χ*^2^ = 0.3451, *p* = 0.3446) and taxa, as fungal taxa abundance comparisons between bat and substrate samples resulted in only one fungal order, *Helotiales*, being more represented in substrate samples than on bats. This result was not caused by the presence of Pd but rather other members of Helotiales, which are common fungi within cave communities.

**Figure 5.**
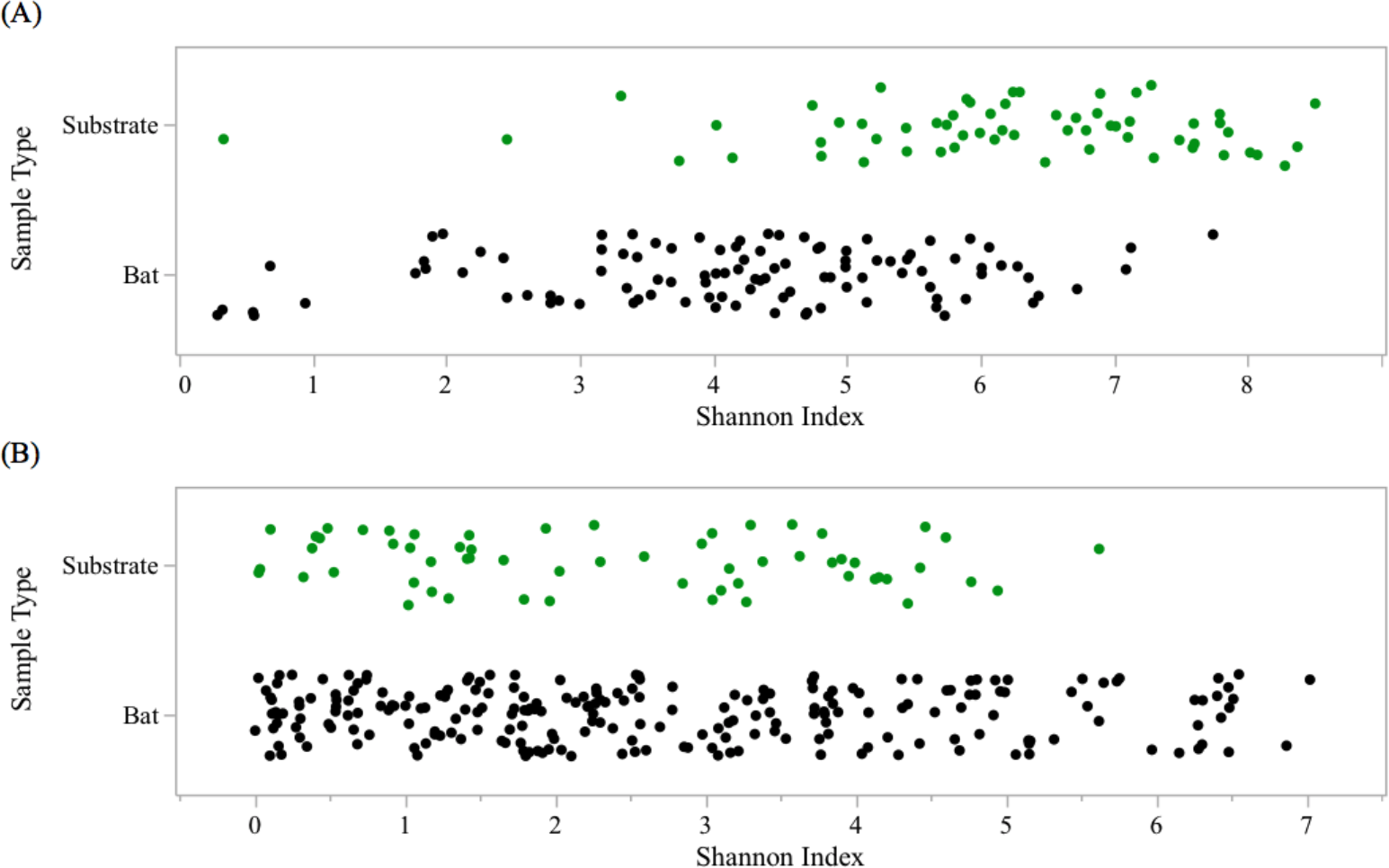
Bacterial (A) and fungal (B) differences between bat epidermal and substrate microbiome composition. Bacterial results indicate a significant difference in diversity between bat and substrate samples, with substrate samples containing a higher Shannon diversity. Fungal results did not indicate a significant difference in diversity between bat and substrate samples.

## Discussion

We found that one of the most heavily impacted species, *M. lucifugus*, which was once highly abundant but underwent massive die-offs from WNS (Frick et al., 2010), is also the bat species whose bacterial microbiome is the most dramatically altered by the presence of Pd. Contrary to expectations, the bacterial microbiome does not appear to have a protective effect but rather is affected after invasion and colonization by Pd, which suggests that Pd results in significant reductions in bacterial community richness and evenness. This result was supported by the modeling, which identified Pd presence as the only significant covariate influencing all three measures of bacterial diversity in *M. lucifugus*. Recent work on host-associated microbial community changes throughout the progression of Sea Star Wasting Disease also found community-wide differences in the microbiomes of affected and unaffected individuals, specifically noting a decrease in species richness of the microbiome in late stages of the disease (Lloyd & Pespeni, 2018). These findings are additionally supported by other wildlife diseases (i.e. chytridiomycosis and snake fungal disease) in which this same effect on host-associated microbial communities are seen (Allender et al., 2018; Jani & Briggs, 2014).

When commensals decrease in abundance throughout the progression of disease, their reduced collective ability to perform functions that inhibit or prevent the growth of pathogens and/or opportunistic bacteria potentially leads to an increase in disease severity. Alternatively, *M. lucifugus* with low richness and evenness may be more susceptible to colonization by Pd. Future work is needed to determine the direction of causation between skin bacterial diversity and disease severity to disentangle these competing hypotheses; however, our findings strongly suggest that Pd invasion causes changes in bat skin microbiomes as data indicates increases in Pd load are correlated with decreases in bacterial diversity.

A bacterial taxonomic differential abundance test between Pd-positive and Pd-negative *M. lucifugus* uncovered several bacterial taxa that are seemingly impacted by the onset and growth of Pd. The *Pseudonocardiaceae* family in the order Actinomycetales was significantly more abundant in Pd-positive bats, while bacterial order *Chromatiales* and bacterial families *Rhizobiaceae* and *Cytophagaceae* were all significantly overrepresented in Pd-negative *M. lucifugus*. Investigations into the *Pseudonocardiaceae* family have found that certain members are involved in the production of antimicrobial agents under specific nitrogen conditions (Platas et al., 1998), with low nitrogen stimulating the production of antibacterial substances in the genera *Amycolatopsis*, *Saccharomonospora*, and *Saccharopolyspora*, and high nitrogen stimulating the production of metabolites in the genus *Pseudonocardia*. Indeed, a well-studied and highly evolved mutualism between fungus-growing ants and their fungi has also uncovered antibiotic-producing bacteria within the *Pseudonocardiaceae* family (Currie, Scott, Summerbell, & Malloch, 1999). Attine ants and their fungi are mutually dependent, with the maintenance of stable fungal monocultures critical to the survival of both organisms. Examination of this symbiotic relationship found that attine ant fungal gardens also host a specialized and virulent parasitic fungus belonging to the genus *Escovopsis* while filamentous bacterium from *Pseudonocardiaceae*—that are largely vertically transmitted between ant generations—produce antibiotics specifically targeted to suppress the growth of the specialized garden parasite (Cafaro & Currie, 2005; Currie, Poulsen, Mendenhall, Boomsma, & Billen, 2006). This relationship in ants suggests a link between members of the *Pseudonocardiaceae* family and fungal pathogen defense, making its overrepresentation in Pd-positive bats interesting as it potentially points to host defense mechanisms.

As for bacterial orders overrepresented in Pd-negative *M. lucifugus*, a key characteristic of the *Cytophagaceae* family is that members of most species are able to degrade one or several kinds of organic macromolecules such as proteins (e.g., casein, gelatin), lipids, starches, and—most noteworthy in this case—chitin (Bergey, 1989). Chitin, a long linear homopolymer of beta-1,4-linked N-acetylglucosamine, is a structurally important component of the fungal cell wall (Bowman & Free, 2006). In both yeasts and filamentous fungi, chitin microfibrils are formed from inter-chain hydrogen bonding. These crystalline polymers significantly contribute to the overall integrity of the cell wall and when chitin synthesis is disrupted, the wall becomes disordered and the fungal cell becomes malformed and osmotically unstable (Bowman & Free, 2006). Given this seemingly important bacterial family characteristic, its abundance in Pd-negative bats suggests a potential link between its enzymatic activity and its ability to limit Pd growth on bat skin. However, we note that this bacterial family was not common in *E. fuscus*, a bat species more tolerant to WNS.

While the *M. lucifugus* bacterial microbiome was impacted by the presence of Pd, *E. fuscus* and *P. subflavus* bacterial microbiomes were similar between the Pd affected and unaffected groups. *Eptesicus fuscus* is one of the least affected species by the introduction of WNS as they still rank among the most abundant and widespread bats in North America (Langwig et al., 2017). In contrast, *P. subflavus* populations appear to be in rapid decline due to WNS (Frick et al., 2017, 2015; Langwig et al., 2012). Thus, it is curious that bacterial diversity in *P. subflavus* was not affected in the same manner as it was in *M. lucifugus*. One potential reason for the difference between *M. lucifugus* and *P. subflavus* has to do with their microbiomes prior to Pd infection. Beta-diversity analyses of Pd-negative *M. lucifugus* and *P. subflavus* (Figs. S3 and S4) indicate their bacterial and fungal microbiomes were significantly different before the onset of disease, and this could potentially explain differences observed following Pd invasion. For fungal community composition, the presence of Pd did not impact fungal taxon richness or evenness in *E. fuscus* or *M. lucifugus*, but did impact the taxon evenness in *P. subflavus*, with significantly higher fungal community evenness in Pd-positive bats. Additionally, our findings suggest an influence of environment on the fungal species found on bats, as there were no significant differences in diversity or taxa between bat skin samples and the cave substrates, a pattern typically associated with transient (non-resident) members of a community (Kong & Segre, 2012; Vanderwolf, Malloch, & McAlpine, 2015). Indeed, the fungal taxa present on bats appear to be a sample of what is present in the environment, with fungal spores adventitiously landing on bat skin rather than a commensal relationship with the fungi living on the host (Holz, Lumsden, Marenda, Browning, & Hufschmid, 2018). Commensal fungi do occasionally grow on bats (e.g. Lorch et al., 2015), but Pd appears to have adapted from an environmental microbe living in cave soils and sediments to a pathogen that is able to utilize bat skin as a food source (Palmer, Drees, Foster, & Lindner, 2018). This was not the case for bacterial species, however, as there were significant differences in diversity between bats and substrates with substrate samples showing higher Shannon diversity. Additionally, relative abundance comparisons resulted in overabundance in more than 20 bacterial families between bat and substrate samples. These results suggest that bat skin serves as a niche for certain bacterial species that remain as commensal members of the bat skin microbiome regardless of environment. Moreover, these commensal bacteria on the bat do not appear to be readily shed into the environment even though the substrate samples were taken in close proximity to the bats.

Inherent complexities in the composition of the microbiome can often preclude investigations of microbe-associated diseases. Instead of single organisms being associated with disease, community characteristics (such as composition and metagenomic functionality) may be more relevant (Cho & Blaser, 2012). Investigations that are able to correlate invasive pathogens and alterations within a microbiome—like the results in this study—potentially suggest that microbiome-host interactions may determine likelihood of infection; however, the contrasting relationship between Pd and the bacterial microbiomes of *M. lucifugus* and *P. subflavus* indicate that we are just beginning to understand how the bat microbiome interacts with a fungal invader such as Pd.

## Supplementary Figure Legends

**Figure S1.**
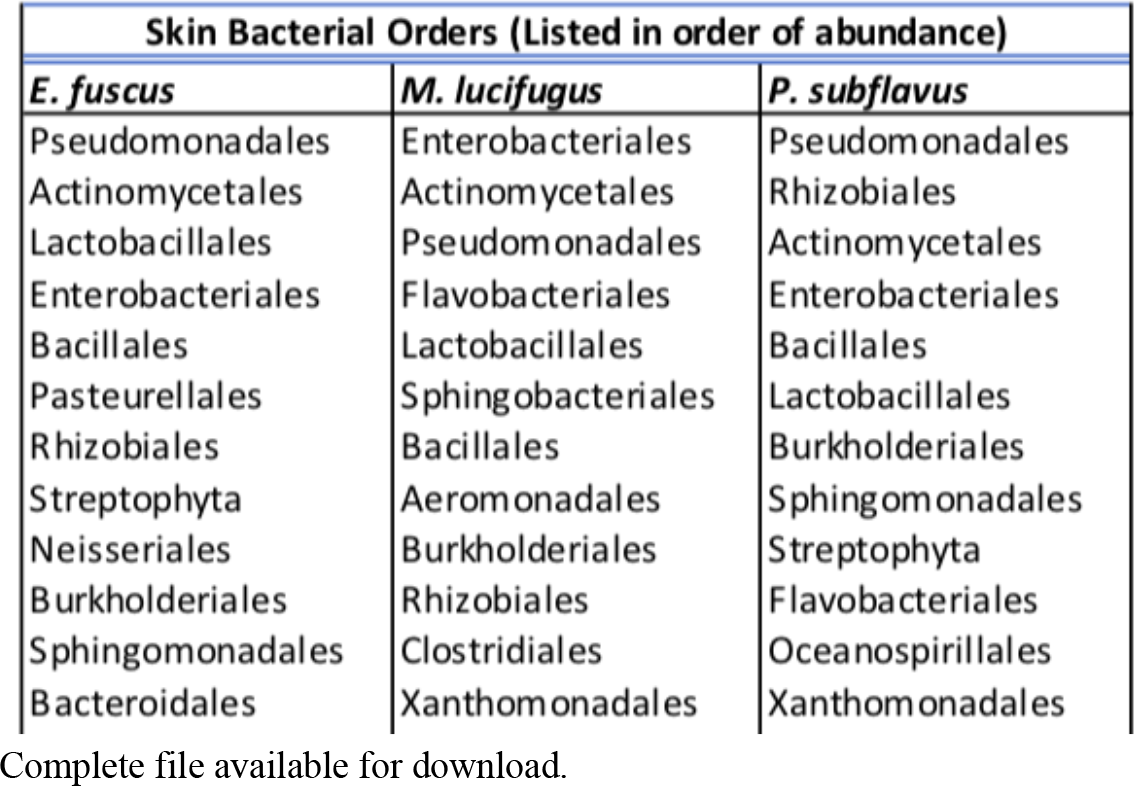
Complete list of skin bacterial orders per species.

**Figure S2.**
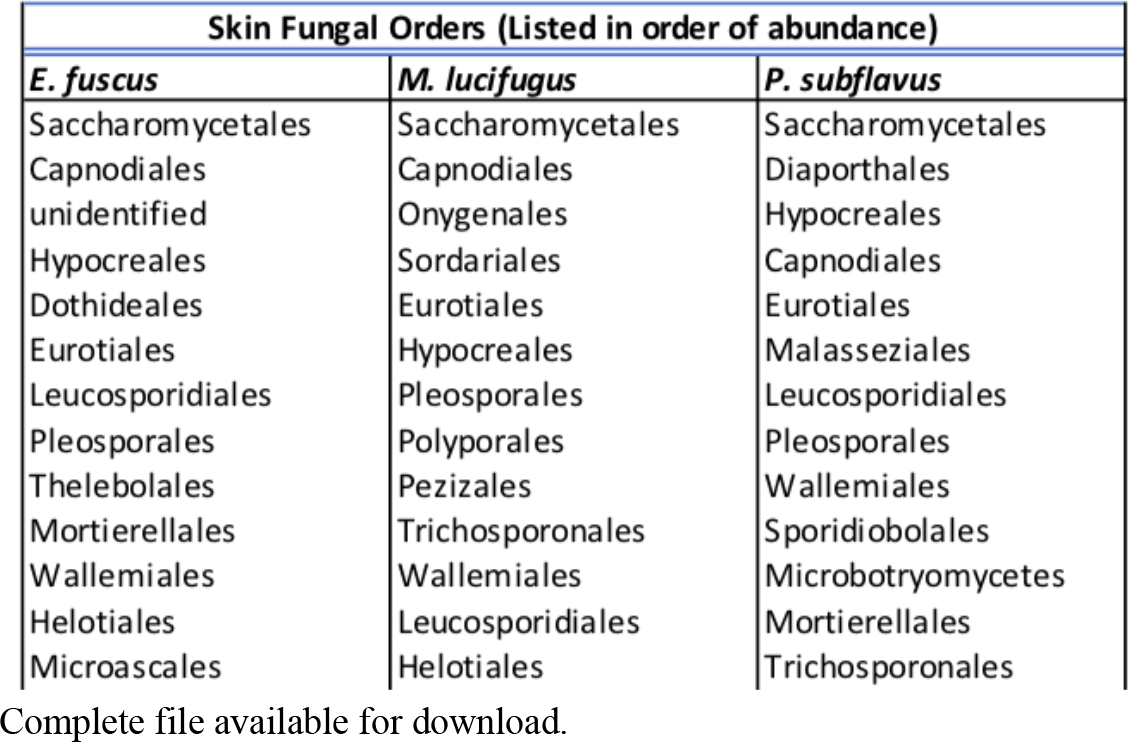
Complete list of skin fungal orders per species.

**Figure S3.**
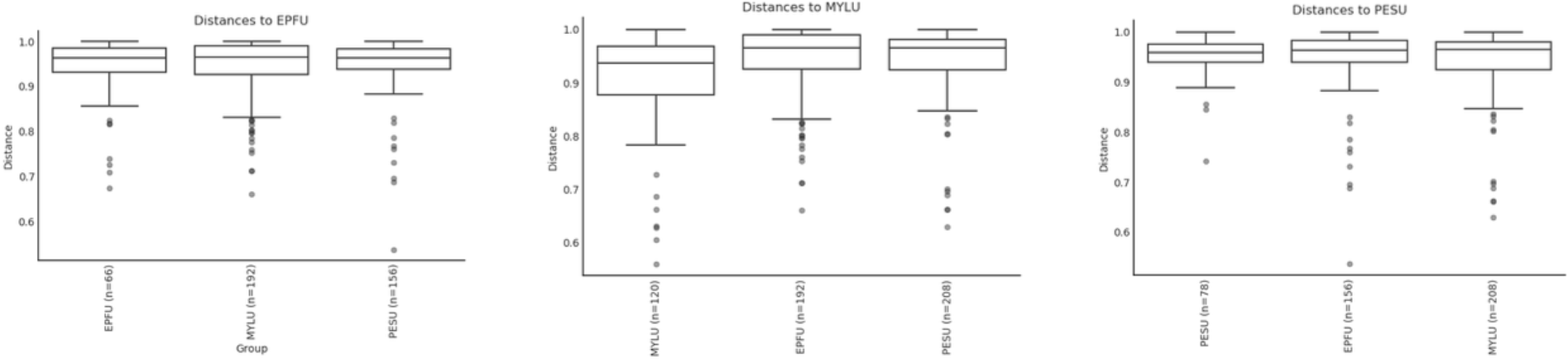
Bacterial beta-diversity analyses between Pd-negative *Eptesicus fuscus*, *Myotis lucifugus*, and *Perimyotis subflavus*. Jaccard distance matrices showed that when measuring bacterial species presence and absence *E. fuscus* and *P. subflavus* were the most similar in bacterial community composition (*p* = 0.19), while *E. fuscus* and *M. lucifugus* were the most dissimilar in bacterial community composition (*p* = 0.01).

**Figure S4.**
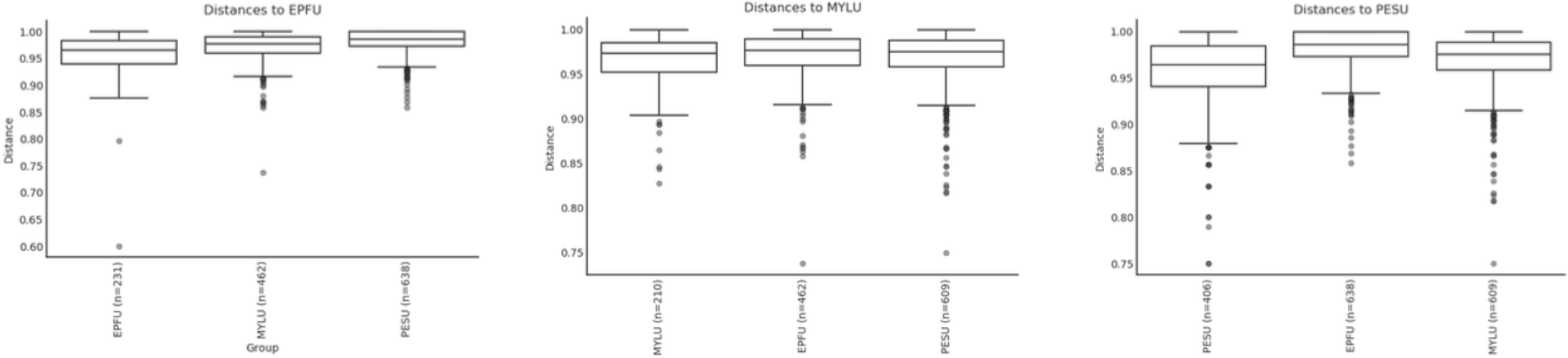
Fungal beta-diversity analyses between Pd-negative *Eptesicus fuscus*, *Myotis. lucifugus*, and *Perimyotis subflavus*. Jaccard distance matrices for fungal community dissimilarity showed high levels of fungal community of dissimilarity between all three species (*E. fuscus* and *M. lucifugus*, *p* = 0.001; *E. fuscus* and *P. subflavus*, *p* = 0.001; *M. lucifugus* and *P. subflavus*, *p* = 0.001).

**Figure S5.**
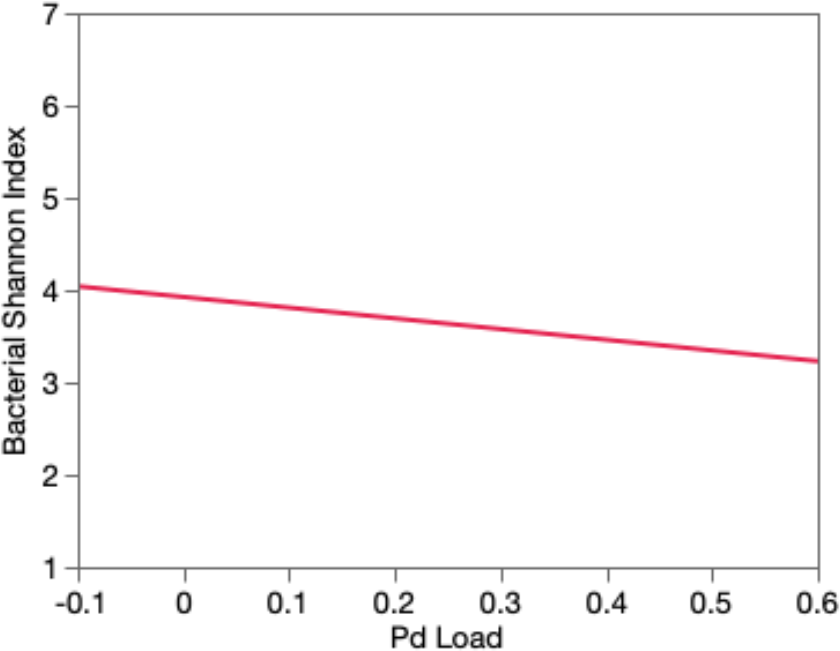
The effect of Pd load on Shannon diversity in *Myotis lucifugus*. Results indicate that increases in Pd load correlated with a decrease in bacterial diversity.

**Figure S6.**
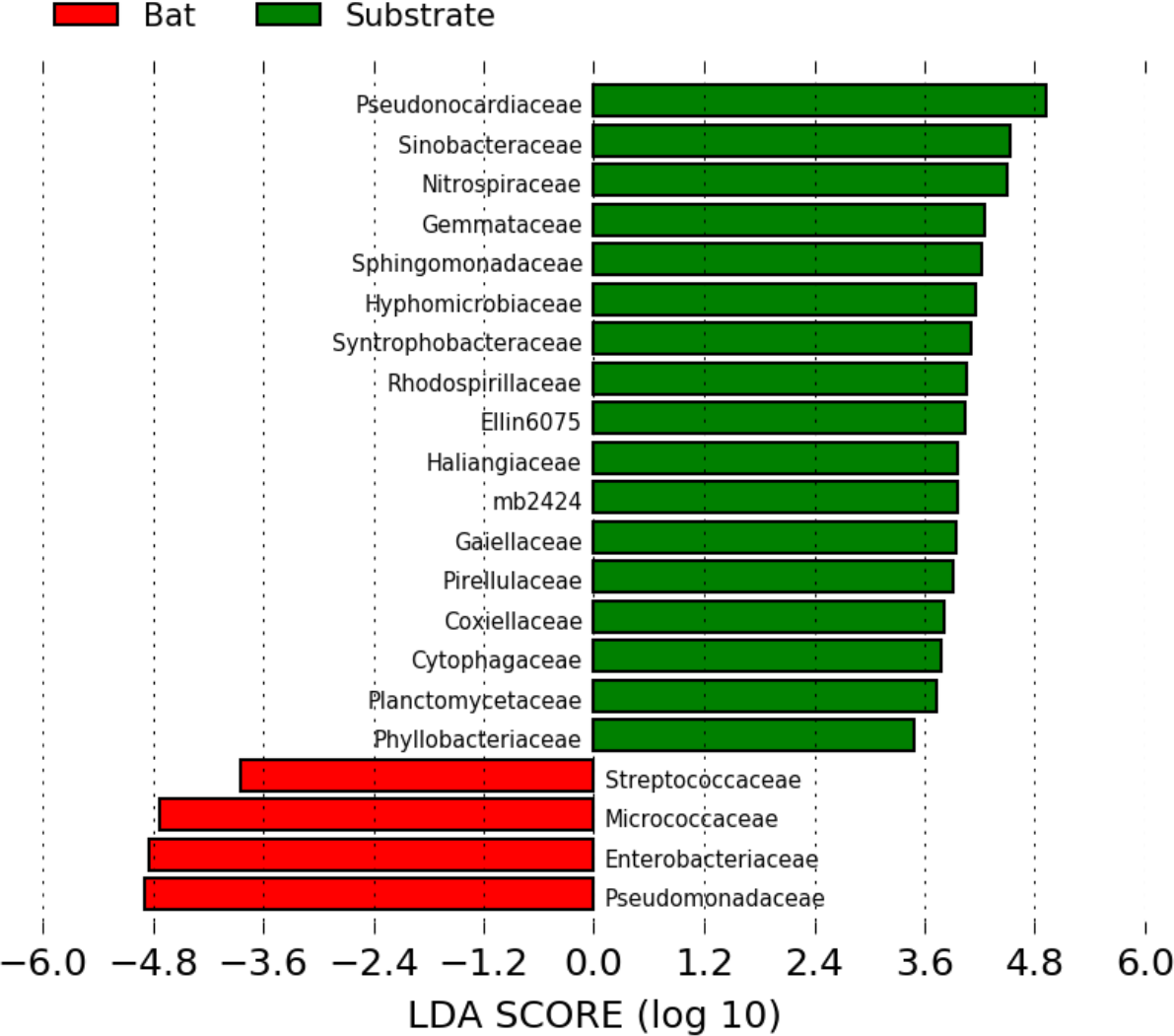
Bacterial differential abundance between bat and substrate samples. Bacterial diversity was significantly higher on substrates than on bats with 17 overly abundant bacterial families represented in the combined substrate samples and only four overly abundant bacterial families represented in combined bat samples

