## Supplemental Figure 1 for "White-nose syndrome restructures bat skin microbiomes"

| Skin Bacterial Orders (Listed in order of abundance) | | |
|---|---|---|
| ***E. fuscus*** | ***M. lucifugus*** | ***P. subflavus*** |
| Pseudomonadales | Enterobacteriales | Pseudomonadales |
| Actinomycetales | Actinomycetales | Rhizobiales |
| Lactobacillales | Pseudomonadales | Actinomycetales |
| Enterobacteriales | Flavobacteriales | Enterobacteriales |
| Bacillales | Lactobacillales | Bacillales |
| Pasteurellales | Sphingobacteriales | Lactobacillales |
| Rhizobiales | Bacillales | Burkholderiales |
| Streptophyta | Aeromonadales | Sphingomonadales |
| Neisseriales | Burkholderiales | Streptophyta |
| Burkholderiales | Rhizobiales | Flavobacteriales |
| Sphingomonadales | Clostridiales | Oceanospirillales |
| Bacteroidales | Xanthomonadales | Xanthomonadales |
| Xanthomonadales | Caulobacterales | Pasteurellales |
| Flavobacteriales | Neisseriales | Chromatiales |
| Sphingobacteriales | Chromatiales | Caulobacterales |
| Saprospirales | Sphingomonadales | Aeromonadales |
| Aeromonadales | Streptophyta | Vibrionales |
| Clostridiales | Myxococcales | Kiloniellales |
| Rhodospirillales | Pasteurellales | envOPS12 |
| Rubrobacterales | Nitrospirales | Saprospirales |
| Oceanospirillales | Saprospirales | Acidimicrobiales |
| Gemellales | Alteromonadales | Rickettsiales |
| Fusobacteriales | iii1-15 | SBR1031 |
| Solirubrobacterales | Solirubrobacterales | Clostridiales |
| Rickettsiales | Fusobacteriales | Alteromonadales |
| Acidobacteriales | RB41 | Campylobacterales |
| Legionellales | Vibrionales | Rhodobacterales |
| Spirochaetales | Bacteroidales | Sphingobacteriales |
| Gemmatales | Syntrophobacterales | Cytophagales |
| RB41 | Cytophagales | Rhodospirillales |
| Alteromonadales | Legionellales | Gemmatales |
| Acidimicrobiales | Bdellovibrionales | Thermales |
| Nitrosomonadales | Gemmatales | Planctomycetales |
| Chromatiales | Nitrosomonadales | Desulfovibrionales |
| Cytophagales | Salinisphaerales | MND1 |
| Caulobacterales | Gemellales | Myxococcales |
| MLE1-12 | Thiotrichales | Thiotrichales |
| DS-18 | Rhodobacterales | iii1-15 |
| Cardiobacteriales | Thermales | agg27 |
| Rhodobacterales | Pedosphaerales | Desulfobacterales |
| Acholeplasmatales | Acidimicrobiales | C114 |
| Thermales | Rickettsiales | Deinococcales |
| Solibacterales | Rhodocyclales | wb1_H11 |

| | | |
|---|---|---|
| Nitrospirales | Gaiellales | Desulfuromonadales |
| Pirellulales | Pirellulales | RB41 |
| Myxococcales | Rhodospirillales | Rhodocyclales |
| Methylophilales | DS-18 | Salinisphaerales |
| Rhodocyclales | Oceanospirillales | pLW-97 |
| Rhodothermales | PK29 | Solirubrobacterales |
| Campylobacterales | Deinococcales | SM2F09 |
| Chlamydiales | Methylophilales | Nitrospirales |
| Ellin329 | Sva0725 | Bacteroidales |
| Gaiellales | Planctomycetales | Neisseriales |
| Syntrophobacterales | Chlorophyta | Rubrobacterales |
| Erysipelotrichales | MIZ46 | Methylophilales |
| Vibrionales | CCU21 | Chthoniobacterales |
| Bdellovibrionales | SM1D11 | Legionellales |
| Mycoplasmatales | NB1-j | Cerasicoccales |
| Thiotrichales | Chroococcales | JG30-KF-CM45 |
| iii1-15 | Campylobacterales | Gemellales |
| Nitriliruptorales | A31 | Pirellulales |
| NB1-j | SJA-36 | Entomoplasmatales |
| CCU21 | envOPS12 | FAC88 |
| PK29 | MND1 | Gemmatimonadales |
| Euzebyales | SBR1031 | mle1-48 |
| JG30-KF-CM45 | Chlamydiales | Erysipelotrichales |
| Desulfovibrionales | PK329 | Opitutales |
| Fimbriimonadales | Kiloniellales | Gaiellales |
| mle1-8 | Solibacterales | BD7-3 |
| Planctomycetales | FAC88 | Chlamydiales |
| Kiloniellales | Methylococcales | Syntrophobacterales |
| SC-I-84 | Rubrobacterales | Verrucomicrobiales |
| MIZ46 | WD2101 | PK29 |
| Sva0725 | Cerasicoccales | Acidobacteriales |
| Phycisphaerales | N1423WL | Sva0725 |
| Deinococcales | Ellin6067 | Spirobacillales |
| AKYG1722 | Verrucomicrobiales | SC-I-84 |
| Chlorophyta | Thiohalorhabdales | Ellin6513 |
| Ellin5290 | Mycoplasmatales | Chroococcales |
| Gemmatimonadales | JG30-KF-CM45 | Oscillatoriales |
| Desulfuromonadales | Entomoplasmatales | Pseudanabaenales |
| Opitutales | Chthoniobacterales | Methylococcales |
| Spirobacillales | Fimbriimonadales | Cryptophyta |
| PHOS-HD29 | agg27 | Pedosphaerales |
| Ignavibacteriales | A21b | Fimbriimonadales |
| Stramenopiles | SC-I-84 | MLE1-12 |
| WCHB1-41 | Bifidobacteriales | Phycisphaerales |
| Pedosphaerales | 258ds10 | |

| | |
|---|---|
| Holophagales | CCM11a |
| 0319-7L14 | Hydrogenophilales |
| Caldilineales | Coriobacteriales |
| SBR1031 | DRC31 |
| Procabacteriales | AKIW781 |
| | Phycisphaerales |
| | PB19 |
| | Cardiobacteriales |
| | Desulfovibrionales |
| | WCHB1-50 |
| | Erysipelotrichales |
| | S0208 |
| | Cryptophyta |
| | Elusimicrobiales |
| | Oscillatoriales |
| | Pseudanabaenales |
| | IS-44 |
| | AKYG885 |
| | d113 |
| | CL500-15 |
