## Supplemental Figure 2 for "White-nose syndrome restructures bat skin microbiomes"

| Skin Fungal Orders (Listed in order of abundance) | | |
|---|---|---|
| ***E. fuscus*** | ***M. lucifugus*** | ***P. subflavus*** |
| Saccharomycetales | Saccharomycetales | Saccharomycetales |
| Capnodiales | Capnodiales | Diaporthales |
| unidentified | Onygenales | Hypocreales |
| Hypocreales | Sordariales | Capnodiales |
| Dothideales | Eurotiales | Eurotiales |
| Eurotiales | Hypocreales | Malasseziales |
| Leucosporidiales | Pleosporales | Leucosporidiales |
| Pleosporales | Polyporales | Pleosporales |
| Thelebolales | Pezizales | Wallemiales |
| Mortierellales | Trichosporonales | Sporidiobolales |
| Wallemiales | Wallemiales | Microbotryomycetes |
| Helotiales | Leucosporidiales | Mortierellales |
| Microascales | Helotiales | Trichosporonales |
| Pezizales | Malasseziales | Helotiales |
| Polyporales | Mortierellales | Polyporales |
| Glomerellales | Thelebolales | Thelebolales |
| Tremellales | Agaricales | Cystofilobasidiales |
| Trichosphaeriales | Dothideales | Russulales |
| Sporidiobolales | Sporidiobolales | Pezizales |
| Malasseziales | Trechisporales | Glomerellales |
| Onygenales | Diaporthales | Microascales |
| Microbotryomycetes | Trichosphaeriales | Dothideales |
| Sordariales | Hymenochaetales | Sordariales |
| Agaricostilbales | Cantharellales | Tremellales |
| Russulales | Russulales | Trichosphaeriales |
| Agaricales | Venturiales | Diversisporales |
| Cantharellales | Tremellales | Agaricales |
| Trichosporonales | Cystofilobasidiales | Hymenochaetales |
| Diversisporales | Chaetothyriales | Onygenales |
| Chaetothyriales | Microascales | Chaetothyriales |
| Cystofilobasidiales | Glomerellales | Trechisporales |
| Hymenochaetales | Microbotryomycetes | Tritirachiales |
| Olpidiales | Xylariales | Xylariales |
| Xylariales | Filobasidiales | Filobasidiales |
| Coniochaetales | Teloschistales | Basidiobolales |
| Ophiostomatales | Spizellomycetales | Amylocorticiales |
| GS11 | Corticiales | Holtermanniales |
| Botryosphaeriales | Taphrinales | Archaeorhizomycetales |
| Filobasidiales | Botryosphaeriales | Taphrinales |
| Archaeorhizomycetales | Togniniales | Mucorales |
| Coryneliales | Olpidiales | Melanosporales |
| Mucorales | Orbiliales | Cantharellales |
| Phacidiales | Agaricostilbales | Olpidiales |

| | | |
|---|---|---|
| Togniniales | Basidiobolales | GS04 |
| GS04 | Auriculariales | Teloschistales |
| Auriculariales | Mucorales | Boletales |
| Trechisporales | Diversisporales | Botryosphaeriales |
| Basidiobolales | Exobasidiales | Phacidiales |
| Teloschistales | GS04 | Septobasidiales |
| Diaporthales | Lichenostigmatales | Calosphaeriales |
| Chaetosphaeriales | GS11 | Ophiostomatales |
| unidentified | Melanosporales | Auriculariales |
| Archaeosporales | Boletales | Lecanorales |
| Venturiales | Ustilaginales | Togniniales |
| Taphrinales | Ophiostomatales | Archaeosporales |
| Amylocorticiales | Atheliales | Agaricostilbales |
| Erysiphales | Peltigerales | GS11 |
| Septobasidiales | Chaetosphaeriales | GS26 |
| Gloeophyllales | Erysiphales | Kriegeriales |
| Boletales | Entylomatales | Coniochaetales |
| Atheliales | Archaeorhizomycetales | Erythrobasidiales |
| Cystobasidiales | Cystobasidiales | Atheliales |
| Exobasidiales | Septobasidiales | Chaetosphaeriales |
| Tritirachiales | Tritirachiales | Dothideomycetes |
| GS34 | Archaeosporales | Venturiales |
| Sebacinales | Lecanorales | Cystobasidiales |
| Rhytismatales | Coniochaetales | Golubeviales |
| | GS26 | |
| | Gloeophyllales | |
| | Entomophthorales | |
| | Dothideomycetes | |
| | Umbilicariales | |
